# Functionally Diverse Coral-Associated Bacteria from an Urban Reef Exhibit Antioxidant and Antibiofilm Properties Relevant to Probiotic Development

**DOI:** 10.1101/2025.07.29.666964

**Authors:** Diana Lorena Gutiérrez-Samacá, Anyela Velasquez-Emiliani, Jordan Ruiz-Toquica

## Abstract

Coral reefs are increasingly threatened by environmental stressors, resulting in substantial ecological losses. Yet, resilient coral reef systems—such as urban reefs—offer a promising reservoir of beneficial microorganisms capable of supporting coral health under stress. In this study, we characterized, and functionally screened bacteria associated with an urban reef dominated by the coral *Madracis auretenra*. From 22 previously selected isolates, all exhibited catalase activity, with several showing notably high levels. Pigmented strains likely produce carotenoids with potential photoprotective and antioxidant functions. Most isolates inhibited biofilm formation in multiple reporter strains. While quorum sensing inhibition, and antagonism against *Vibrio coralliilyticus* were observed, both effects were generally low, suggesting that suppression of pathogens may occur primarily through interference of biofilm formation rather than direct inhibition. Growth kinetics revealed strain-specific dynamics, with fast-growing isolates offering potential advantages for probiotic application. Multidimensional functional profiling indicated trait overlap and redundancy among strains, reflecting niche specialization and the co-expression of multiple beneficial traits—features desirable for the design of stable probiotic consortia. These findings underscore the probiotic potential of bacteria associated with urban *M. auretenra*, particularly for traits related to oxidative stress mitigation and microbial regulation. Further *in vivo* validation is needed to assess their efficacy under reef-relevant stress conditions.

## Introduction

Coral reefs are under increasing pressure from both environmental and anthropogenic stressors, with global coral cover having declined by 50% since the 1950s, leading to significant losses in biodiversity and structural complexity (Alvarez-Filip et al., 2009; Eddy et al., 2021). In this context, urban coral reefs represent a valuable study model due to the challenging environments they inhabit, characterized by high turbidity, pollution, and eutrophication associated with coastal urban development (Burt et al., 2020; Dsikowitzky et al., 2016; Heery et al., 2018; Zweifler et al., 2021). These coral assemblages have developed acclimatization and adaptation mechanisms, including symbiotic interactions with bacteria that enable them to persist in such altered environments (Lima et al., 2023; Wainwright et al., 2019).

This resilience, driven by sustained environmental stress, is thought to result from dynamic adjustments in the coral microbiome (Ruiz-Toquica et al., 2025; Ziegler et al., 2019). These adjustments promote flexible host–microbe associations that optimize microbial functional traits to counteract prevailing stressors, thereby enhancing coral health, survival, and ecosystem persistence (Röthig et al., 2020). In this framework, the identification and application of beneficial microorganisms for corals—commonly referred to as coral probiotics (Peixoto et al., 2021)—has emerged as a promising strategy to manipulate the microbiome, reverse dysbiosis, and restore holobiont functionality (Peixoto et al., 2017; Voolstra & Ziegler, 2020).

Multiple studies have demonstrated the potential of beneficial coral bacteria to mitigate stress under laboratory conditions, including contaminant degradation, enhanced thermal tolerance, and recovery from experimental bleaching (Cardoso et al., 2024; Fragoso ados Santos et al., 2015; Morgans et al., 2020; Santoro et al., 2021). Functional screening of candidate strains typically focuses on traits such as antioxidant capacity, antimicrobial activity, and microbial signaling disruption—features associated with increased coral resilience (de Breuyn et al., 2025; Peixoto et al., 2021).

Among these, microbial antioxidant activity is considered a key mechanism for mitigating thermal stress, as it reduces reactive oxygen species (ROS) that accumulate during bleaching events (Dungan et al., 2021; Li et al., 2023; Rosado et al., 2019). Enzymes such as catalase, peroxidase, and superoxide dismutase have been identified in coral-associated bacteria, including those from *Galaxea fascicularis*, and successfully applied as probiotics in *Mussismilia hispida* and *Pocillopora damicornis*, enhancing resistance to bleaching and recovery under heat stress (Doering et al., 2023; Rosado et al., 2019; Santoro et al., 2021).

In addition to enzymatic defenses, photoprotective compounds such as carotenoids and mycosporine-like amino acids (MAAs) also contribute to ROS neutralization and UV protection. These natural “sunscreens” have been reported in microbial symbionts of *Porites lobata* and *Acropora gemmifera*, demonstrating their role in protecting the coral holobiont from excessive solar radiation (Dunlap & Shick, 1998; Kirilovsky & Kerfeld, 2012; Palma Esposito et al., 2025).

Coral-associated bacteria also produce antimicrobial compounds that help structure and regulate microbial communities. These traits are essential for protecting corals from opportunistic infections and disease outbreaks, particularly when host immunity is compromised by prolonged stress (Kvennefors et al., 2012; Mascuch et al., 2023; Ritchie, 2006). The effectiveness of probiotic-derived compounds has been demonstrated against *Vibrio coralliilyticus*, as well as in the control of diseases such as Stony Coral Tissue Loss Disease (SCTLD) and white pox caused by *Serratia marcescens* (Irudayarajan et al., 2024; Monti et al., 2024; Pitts et al., 2025; Reina et al., 2022; Ushijima et al., 2023).

Another promising probiotic trait is the ability to disrupt quorum sensing (QS), a bacterial cell–cell communication system that regulates virulence, antibiotic production, and biofilm formation (Sharp & Ritchie, 2012). Several coral diseases are linked to QS-regulated processes, including bleaching and tissue degradation driven by opportunistic pathogens via N-acyl homoserine lactone (AHL) signals (Certner & Vollmer, 2015; Zhou et al., 2020). The use of quorum sensing inhibitors provides a microbial strategy for controlling pathogenic colonization and community-level imbalances (Bhardwaj et al., 2013; Fetzner, 2015; Xu et al., 2023). This has been explored in the context of coral diseases such as white band, caused by *Vibrio carchariae* and *V. harveyi*, and black band, associated with polymicrobial consortia (Certner & Vollmer, 2015; Meyer et al., 2016; Ritchie & Smith, 1998; Zimmer et al., 2014).

Despite growing interest in urban corals as models for microbial resilience, little is known about the functional potential of their associated bacteria as sources of beneficial symbionts. While preliminary studies have identified their microbiome stability under stress (Ruiz-Toquica et al., 2025; Ziegler et al., 2019), their utility for probiotic development remains largely unexplored. In this study, we characterized, and functionally screened bacteria associated with an urban reef dominated by the coral *Madracis auretenra*, aiming to identify candidate strains with traits relevant to coral health—specifically antioxidant production, quorum sensing disruption, and biofilm inhibition. These traits were used to inform the rational selection of beneficial microbes for potential use in probiotic formulations designed to enhance coral resilience.

## Methods

### Bacterial strains

The bacterial isolates evaluated in this study (Table S1) were previously obtained from the mucus and tissue of urban *Madracis auretenra* corals (Ruiz-Toquica et al., 2023). Based on initial qualitative screening for probiotic potential, 32 isolates were selected out of 132 for detailed analysis and are hereafter referred to as tester strains. These isolates were cryopreserved in liquid media supplemented with 20% glycerol and stored at –20 °C for up to one year.

Target strains included the coral pathogen *Vibrio coralliilyticus* (kindly provided by Kimberly Ritchie, University of South Carolina Beaufort), as well as reference strains *Pseudomonas aeruginosa* ATCC® 27853™, *Escherichia coli* ATCC® 25922™, and *Staphylococcus aureus* ATCC® 25923™, obtained from the Water Quality Laboratory at the University of Magdalena. Additionally, *Chromobacterium violaceum* and *Bacillus subtilis* strains were supplied by the “Study and Utilization of Marine Natural Products and Fruits of Colombia” research group at the National University of Colombia.

### Phylogenetic analysis of *Vibrio* isolates

Due to 24 out of the 32 tester strains were affiliated to *Vibrio*, a genus that typically includes several known coral pathogens, we performed a phylogenetic analysis to evaluate their proximity to pathogenic strains. We compiled 16S rRNA gene sequences from *Vibrio* strains isolated from healthy and diseased corals, as reported by Sweet et al. (2021), and included the sequences obtained in this study (Table S2). Multiple sequence alignment was conducted using MUSCLE, followed by phylogenetic tree reconstruction employing the neighbor-joining method with default parameters and 1,000 bootstrap replicates in MEGA X (v11.0.13). The 16S rRNA gene from *Thermotoga maritima* MSB8 served as an outgroup, reflecting its basal phylogenetic position within Bacteria. The resulting tree was visualized using the *ggtree* package in R (v4.2.2). Strains clustering closely with known pathogens or sequences from diseased coral tissues were excluded from subsequent analyses to focus on putative non-pathogenic candidates.

### Revival and growth conditions

Tester strains were revived and maintained by biweekly subculturing on Tryptic Soy Agar (TSA, Condalab, Spain) supplemented with 1% NaCl. Assays employed various media, including Luria-Bertani (LB) broth and agar (10 g/L peptone, 10 g/L NaCl, 5 g/L yeast extract, with 15 g/L agar when needed), Marine Agar (MA, Condalab), Tryptic Soy Broth (TSB, Condalab), and phosphate-buffered saline (PBS; 137 mM NaCl, 2.7 mM KCl, 100 mM Na_2_HPO_4_, 2 mM KH_2_PO_4_, pH 7.4). *C. violaceum* was maintained on TSA without NaCl. Cultures were incubated overnight at 28°C, with liquid cultures shaken at 145 rpm.

### Extracellular products (ECPs) preparation

ECPs from the tester strains were prepared for liquid assays following a modified protocol based on Cabo et al. (1999) and Villamil et al. (2010). Overnight cultures were diluted in PBS to an optical density at 600 nm (OD_600_) of 0.4–0.6 (Zhang et al., 2015) using a Modulus™ Multimode Microplate Reader (Turner BioSystems, USA). Diluted cultures were inoculated into 10 mL LB broth and incubated with agitation for 72 h. Cultures were centrifuged at 13,000 rpm for 10 min, and the supernatant was filtered through a 0.22 µm membrane (Millipore). Sterile ECP-containing supernatants were stored at –20°C until use (Sheng et al., 2005).

### Catalase activity assay

Catalase (CAT) activity was measured to assess antioxidant potential, following Aebi (1984) with modifications. The reaction mixture included 680 µL of 100 mM phosphate buffer (pH 7.0) containing 10 mM H_2_O_2_ and 1.32 mL of ECPs. The H_2_O_2_ decomposition was monitored at 240 nm, and CAT activity was calculated based on the decrease in absorbance per minute. One unit (U) was defined as the amount of enzyme degrading 1 µM H_2_O_2_ per minute, normalized to protein content (U mg^−1^ protein) (Anithajothi et al., 2014). ECPs from *S. aureus* and *E. coli* served as a positive controls and sterile LB medium was used as a negative control.

### Pigment production analysis

Several bacterial isolates exhibited visible pigment production (Ruiz-Toquica et al., 2023; Table S1). To characterize pigment types and estimate yield, pigments were extracted from 100 mL cultures grown for 72 h (initial OD_600_ = 0.1–0.2). Cultures were mixed 1:1 with 100% methanol and centrifuged at 3,000 rpm for 15 min to obtain the colored supernatant. Absorbance spectra of the extracts were recorded at 25 nm intervals from 400 to 700 nm using a GENESYS 10 UV spectrophotometer, and pigment identity was inferred from spectral peaks (Ratnakaran et al. 2020). Pigment yield was estimated by filtering the extracts through Whatman® grade 1 filter paper, drying the retained material in a muffle furnace at 60°C, and calculating dry weight (mg mL⁻¹) relative to the initial extract volume.

For chromatographic profiling, three isolates that produced sufficient pigment were selected. Fresh extracts (10 mg mL⁻¹ in 100% HPLC-grade methanol) were obtained from wet biomass, sonicated for 30 min at 25°C in the dark, and filtered through 0.22 µm PTFE membranes. Pigments were analyzed using a Thermo Scientific Dionex UltiMate 3000 UHPLC system equipped with a diode array detector (DAD) and evaporative light scattering detector (ELSD). Separation was performed using a LiChroCART® RP-C18e column (5 µm, 4.6 × 250 mm) under the following gradient: 0–5 min (90% methanol), 8–18 min (100% methanol), 21–26 min (90% methanol), at a constant flow rate of 1.0 mL min⁻¹. DAD spectra were recorded from 200 to 800 nm. All chromatographic analyses were conducted at the Chemistry Laboratory of the National University of Colombia.

### Biofilm inhibition

The biofilm inhibition potential of tester strains was assessed following O’Toole (2011). ECPs (20 µL) were added to 96-well plates containing 180 µL of LB cultures (OD_600_ = 0.1–0.2) of four biofilm-forming target strains: *P. aeruginosa*, *E. coli*, *B. subtilis*, and *V. coralliilyticus*. Plates were incubated for 24 h at 28 °C.

After incubation, wells were washed with deionized water, air-dried (10 min), fixed with 250 µL methanol (15 min), and dried again. Biofilms were stained with 1% crystal violet (200 µL, 20 min), washed three times, and air-dried (15 min). Stain was solubilized with 250 µL of 95% ethanol, and absorbance was read at 600 nm. LB broth served as a negative control. Assays were performed in triplicate, and biofilm inhibition was calculated as: % biofilm inhibition = ((Blank OD_600_ – Tester OD_600_) /Blank OD_600_) × 100.

### Inhibition of quorum sensing in *Chromobacterium violaceum*

Quorum sensing (QS) inhibition was evaluated using *C. violaceum*, which produces the pigment violacein under QS control (Reina et al., 2019). In the qualitative assay, *C. violaceum* was spread on LB agar plates, and sterile 5 mm paper discs were placed on the surface. Each disc was inoculated with 3 µL of tester strain culture (OD_600_ = 0.1–0.2 in PBS). Plates were incubated for 48 h at 28 °C, and QS inhibition was indicated by a colorless halo around the discs. LB broth served as the negative control and assays were performed in triplicate.

Strains that exhibited quorum sensing inhibition in the agar-based assay were further evaluated using a quantitative liquid assay in 96-well plates. Each well was inoculated with 20 µL of tester strain ECPs and 180 µL of *C. violaceum* culture (OD_600_ = 0.1–0.2 in LB medium). As pigment extraction was not conducted in this study to quantify violacein at 585 nm, absorbance was measured at 600 nm to assess total growth and pigment production in the wells after 48 h of incubation. Percent inhibition was calculated as: % violacein inhibition = ((Blank OD_600_ – Tester OD_600_) /Blank OD_600_) × 100.

### Inhibition of Vibrio coralliilyticus

The antagonistic activity of tester strains against *V. coralliilyticus*—a well-known coral pathogen also linked to coral bleaching—was assessed via disk diffusion (Reina et al., 2019). *V. coralliilyticus* was spread on LB agar plates, and sterile paper discs were loaded with 10 µL of each tester isolate (OD_600_ = 0.1–0.2 grown in TSB + 1% NaCl). Inhibition was recorded as a clear halo surrounding the discs. Sterile TSB served as the negative control and assays were performed in triplicate.

For isolates showing activity, inhibition was quantified in 96-well microplates following the same protocol used for *C. violaceum*. Wells received 20 µL of tester ECPs and 180 µL of *V. coralliilyticus* culture (OD_600_ = 0.2). After 48 h incubation, absorbance at 600 nm was measured, and inhibition was calculated as: % inhibition of *V. coralliilyticus* = ((Blank OD_600_ – Tester OD_600_) /Blank OD_600_) × 100.

### Growth curves

Isolates exhibiting high activity in at least one of the evaluated traits, were selected for growth kinetics analysis following Villela et al. (2019). Overnight cultures were adjusted to an initial OD_600_ of 0.1–0.2, and 100 µL was inoculated into 10 mL of TSB supplemented with 1% NaCl. Cultures were incubated at 28 °C with agitation (145 rpm), and OD_600_ was measured at 0, 6, 12, 24, 30, 36, 48, 60, and 72 h using a 96-well plate format. All measurements were performed in triplicate. Growth curves were constructed by plotting OD_600_ against time.

### Statistical analyses

All assays were conducted in triplicate. Data were first tested for normality using the Shapiro–Wilk test and for homogeneity of variance using Levene’s test. When assumptions of normality or equal variance were not met, equivalent non-parametric tests were applied. Catalase activity was analyzed via one-way ANOVA to assess differences among tester strains by coral compartment (mucus and tissue) and compared to controls. Biofilm inhibition was evaluated for each reporter strain using the Kruskal-Wallis test followed by Dunn’s post hoc test with Bonferroni correction. Inhibition of QS activity was assessed using the Mann-Whitney U test. Differences in *V. coralliilyticus* inhibition by isolate source (mucus or tissue) were analyzed using a two-tailed Welch’s t-test. Additionally, the area under the curve (AUC) was calculated from growth curves to compare growth performance between bacterial isolates.

Probiotic activity profiles were compared using permutational multivariate analysis of variance (PERMANOVA; Anderson, 2001), based on a Bray–Curtis similarity matrix constructed from untransformed data. The analysis employed Type III sums of squares with 9,999 permutations, and pairwise differences were assessed using Monte Carlo *p*-values in PRIMER6. To identify which isolates contributed most to the observed differences in activity, a Similarity Percentage (SIMPER) analysis was conducted (Anderson & Robinson, 2003). Patterns in functional trait variation among bacterial isolates from the mucus and tissue of coral *Madracis auretenra* were visualized using non-metric multidimensional scaling (NMDS), performed in RStudio (v4.3.3) using the same Bray–Curtis matrix.

## Results

### Phylogenetic relationship of *Vibrio* isolates

Seven bacterial isolates out of 24 strains affiliated with *Vibrio* from the present work were excluded based on their phylogenetic proximity to known coral pathogens or to strains previously recovered from diseased coral tissues (Fig. 1). This refinement yielded a final set of 22 bacterial isolates (Table S3) selected for quantitative assessment of probiotic traits relevant to coral health. Additionally, isolates ICT35, ICM64, and PCNM4 were excluded due to inconsistent growth during initial culturing, precluding their inclusion in downstream analyses.

**Figure 1.**
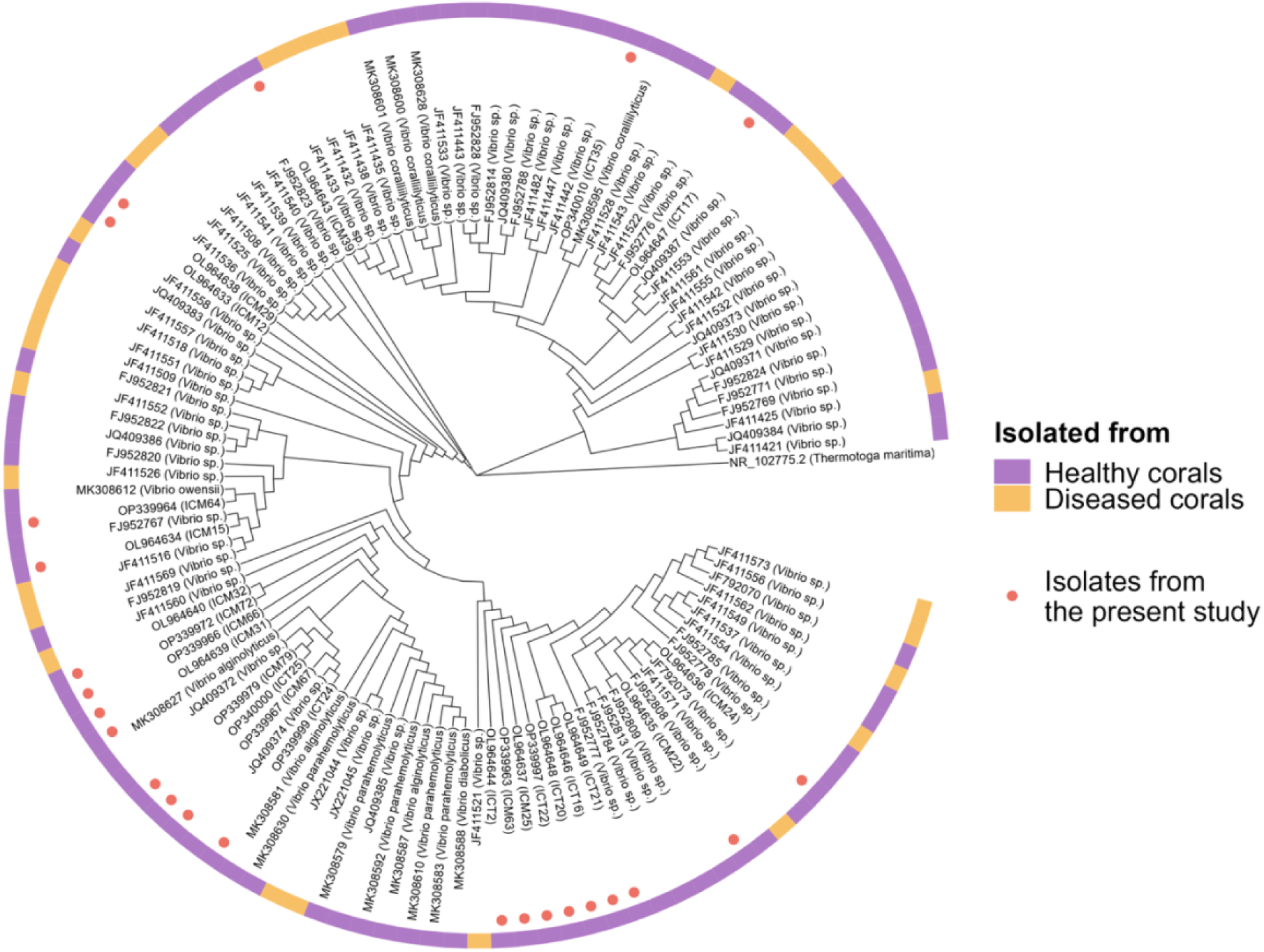
Phylogenetic relationships of *Vibrio* isolates from urban *Madracis auretenra* in the context of known *Vibrio* spp. from healthy and diseased corals. Maximum likelihood tree based on 16S rRNA gene sequences showing the phylogenetic placement of *Vibrio* isolates obtained in the present study (red circles) relative to reference sequences from healthy (purple bars) and diseased (orange bars) coral hosts. The tree was rooted using *Thermotoga maritima* as the outgroup. Reference sequences were retrieved from Sweet et al. (2021) and include representatives from diverse coral genera and health states. Colored bars indicate the health status of the coral source from which each reference strain was isolated. This analysis informed the exclusion of isolates phylogenetically close to potential pathogens or opportunists and guided the selection of strains for subsequent screening.

### Antioxidant and antibiofilm properties of coral bacteria

All 22 bacterial isolates exhibited catalase activity, with values ranging from 1.0 to 6.0 U mg⁻¹ protein (Fig. 2A). No significant differences were observed among isolates based on their source of isolation when compared to controls (One-way ANOVA; *F* _(3, 68)_ = 1.85; *p =* 0.15). Notably, isolates ICM72, ICT22, ICM12, and ICM37—affiliated with the genera *Vibrio* and *Shewanella*—showed catalase activities equal to or exceeding that of the positive control, *Staphylococcus aureus* (6.1 U mg⁻¹ protein).

**Figure 2.**
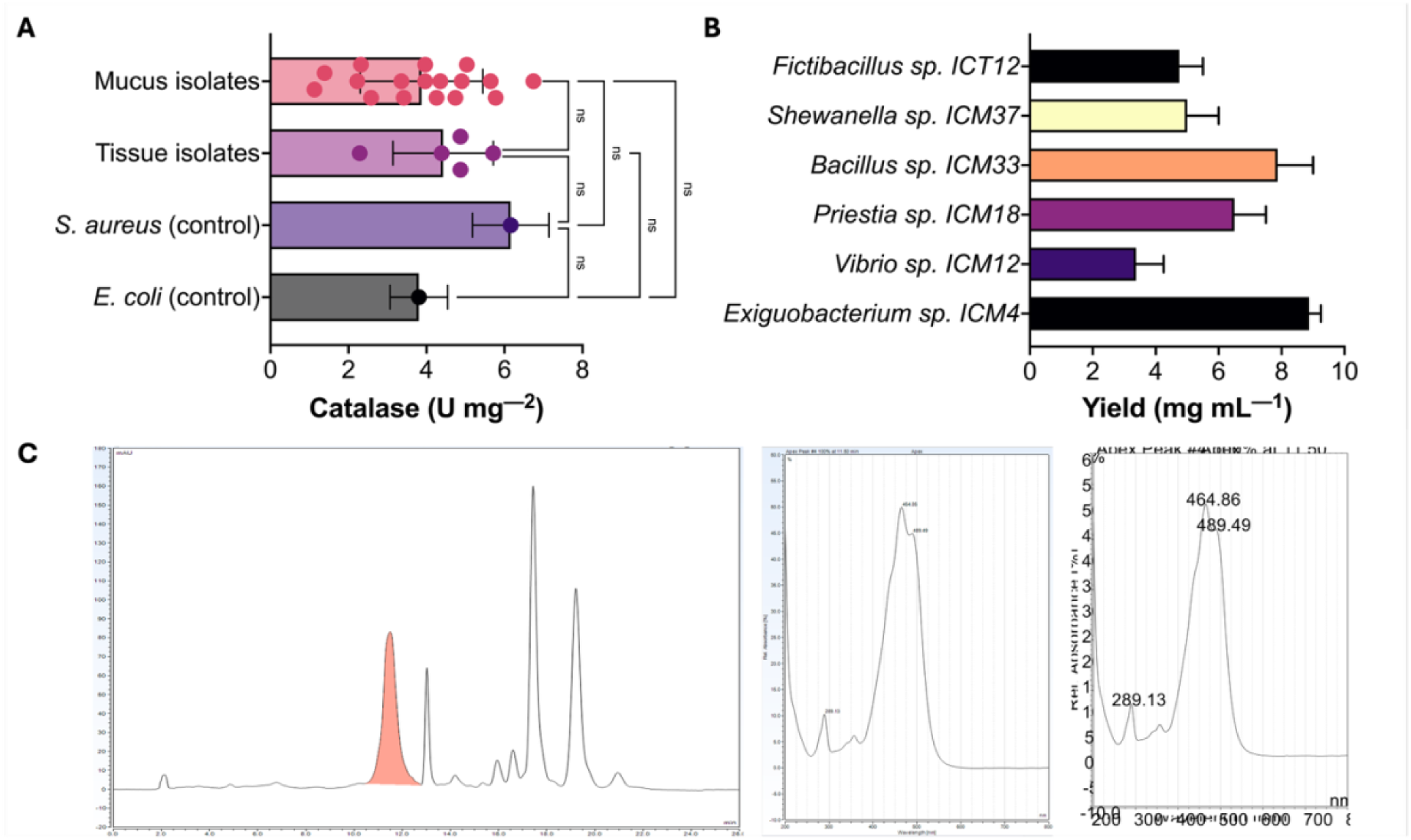
Antioxidant and pigment properties of bacteria isolated from the urban coral *Madracis auretenra*. **(A)** Catalase (CAT) activity of bacterial isolates measured in U mg^−1^ protein. Isolates were grouped by coral compartment of origin (mucus or tissue). Values for the positive controls (*Staphylococcus aureus* and *Escherichia coli*) are also shown. Individual data points represent mean values of the single strains, with bars showing overall mean ± standard deviation. Some isolates performed equal or better than the positive controls (>4 U mg^−1^). **(B)** Pigment yield (mg mL^−1^) of selected isolates that produced colored pigments. Absorbance spectra were consistent with carotenoid-like compounds, which are known to have antioxidant and photoprotective properties. **(C)** Representative chromatogram of a methanolic extract from *Exiguobacterium sp.* ICM4, showing multiple peaks in the 10–20 min range. The highlighted peak (11–13 min) corresponds to a dominant carotenoid-like compound detected across several pigment-producing isolates.

Six isolates not previously excluded from the analysis exhibited pigment production (Table S1). Their absorbance spectra showed peaks ranging from 400 to 500 nm (Fig. S1), consistent with carotenoid-type pigments. Pigment concentrations ranged from 4 to 10 mg mL⁻¹, with *Exiguobacterium* sp. ICM4 producing the highest yield (Fig. 2B).

Chromatographic profiling of extracts from three selected isolates revealed distinct peaks between 10–20 min (Fig. 2C; Fig. S2), indicative of low-to non-polar compounds. A dominant peak observed across all chromatograms, spanning 11–19 min (highlighted in red), further supports the presence of carotenoid-like molecules. UV-visible absorbance patterns revealed maxima between 444 and 493 nm, characteristic of hydrophobic pigments with conjugated double bonds such as zeaxanthin, β-carotene, lycopene, or astaxanthin (Table S4). While definitive identification requires further structural analysis, the spectral and chromatographic signatures strongly suggest the production of carotenoids with potential photoprotective and antioxidant functions by coral-associated bacteria inhabiting urban reef environments.

Most isolates inhibited biofilm formation by more than 40% (Fig. 3A). Significant differences in inhibition activity were observed among the isolates for each reporter strain (Kruskal-Wallis; *H* _(1, 3)_ = 185.33; *p* < 0.001, Table S5). Several candidate strains were particularly effective against biofilms formed by *Escherichia coli*, *Pseudomonas aeruginosa*, and *Bacillus subtilis*, achieving inhibition levels above 75%. In contrast, inhibition of *Vibrio coralliilyticus* biofilms ranged from 12% to 42%, with extracellular products (ECPs) from isolates *Exiguobacterium* sp. ICM4, *Vibrio* sp. ICM67, and *Bacillus* sp. PCNM6 showing the highest activity (Fig. 3A).

**Figure 3.**
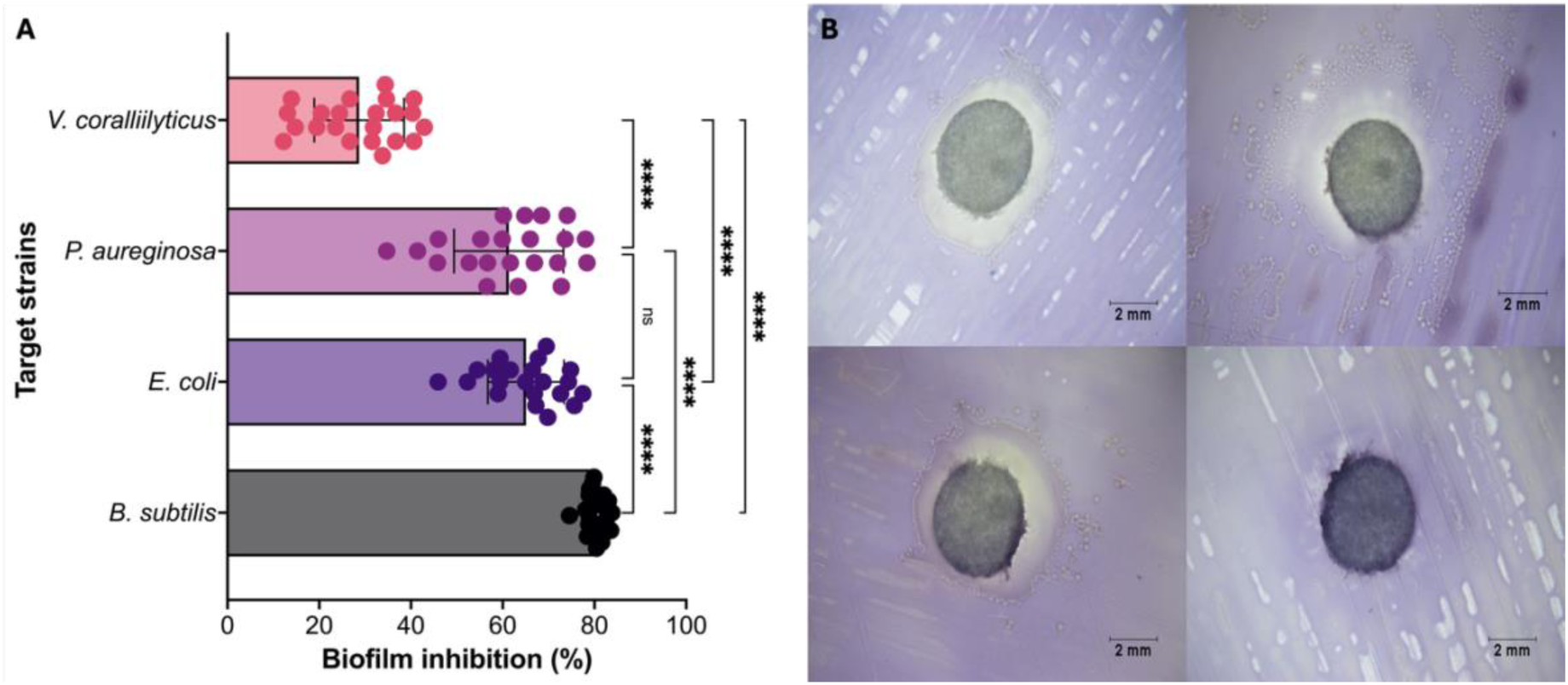
Biofilm inhibition by bacterial isolates from the urban coral *Madracis auretenra*. **(A)** Quantitative assessment of biofilm inhibition against four target strains: *Vibrio coralliilyticus*, *Pseudomonas aeruginosa*, *Escherichia coli*, and *Bacillus subtilis*. Each point represents the mean biofilm inhibition percentage value of a single coral isolate against a specific target. Bars indicate overall mean ± standard deviation. Asterisks indicate significant differences among treatments based on Kruskal-Wallis and Dunn’s post hoc test (\*\*\*\**p* < 0.0001; \*\*\**p* < 0.001; ns = not significant). **(B)** Representative micrographs of violacein inhibition halos surrounding bacterial colonies of coral isolates, indicating varying degrees of quorum sensing disruption in the reporter strain *Chromobacterium violaceum*. Bottom right image shows the negative control (sterile medium with no inhibition). Scale bars = 2 mm.

### QS inhibition, antagonism and growth performance

Most isolates inhibited violacein production by *Chromobacterium violaceum* in agar-based assays, as evidenced by reduced pigment production compared to the sterile medium control (Fig. 3B). While some isolates maintained this activity in the liquid assay, their inhibitory effect was generally low (< 40%) (Fig. S3). Notably, inhibition levels varied depending on the isolate’s source of origin, with differences observed between tissue- and mucus-derived strains (Mann-Whitney; U = 96; *p* = 0.04, Table S5). Notably, isolates *Bacillus* sp. PCNM6 and *Fictibacillus* sp. ICT12 exhibited the highest inhibition of violacein production, ranging from 34% to 36% (Fig. S3).

Despite several isolates showing antibiofilm activity against *V. coralliilyticus*, their ability to directly inhibit the growth of this pathogen was markedly lower. No significant differences were detected between isolates based on their source of isolation (*t* _(15.72)_ = –1.17; *p* = 0.26, Table S5). Inhibition was minimal in both the agar-based assay and the liquid assay, with growth suppression ranging from 5% to 14% (Fig. S4). Among them, isolates *Vibrio* sp. ICM79 and *Fictibacillus* ICT12 demonstrated the most consistent antagonistic activity against *V. coralliilyticus*.

Growth performance was successfully assessed for 14 bacterial isolates. Most exhibited comparable growth kinetics (Fig. 4A). *Exiguobacterium* sp. ICM4, *Bacillus* sp. ICM18, and *Fictibacillus* sp. ICT12 demonstrated rapid growth and superior performance, reaching the highest OD_600_ values between 48 and 72 h, as supported by their area under the curve (AUC) values (Fig. S5). Isolates affiliated with *Vibrio* showed accelerated growth within the first 12 h, followed by a stable stationary phase up to 48 h. In contrast, *Bacillus* sp. ICM33 and *Shewanella* sp. ICM37 exhibited slower growth rates, yet their AUC values indicated moderate overall performance. Notably, *Vibrio* isolates ICM15 and ICM12 grew faster initially but yielded lower AUC values, reflecting reduced cumulative growth (Fig. 4A and Fig. S5). These results highlight strain-specific growth dynamics, which may influence their potential application as coral probiotics.

**Figure 4.**
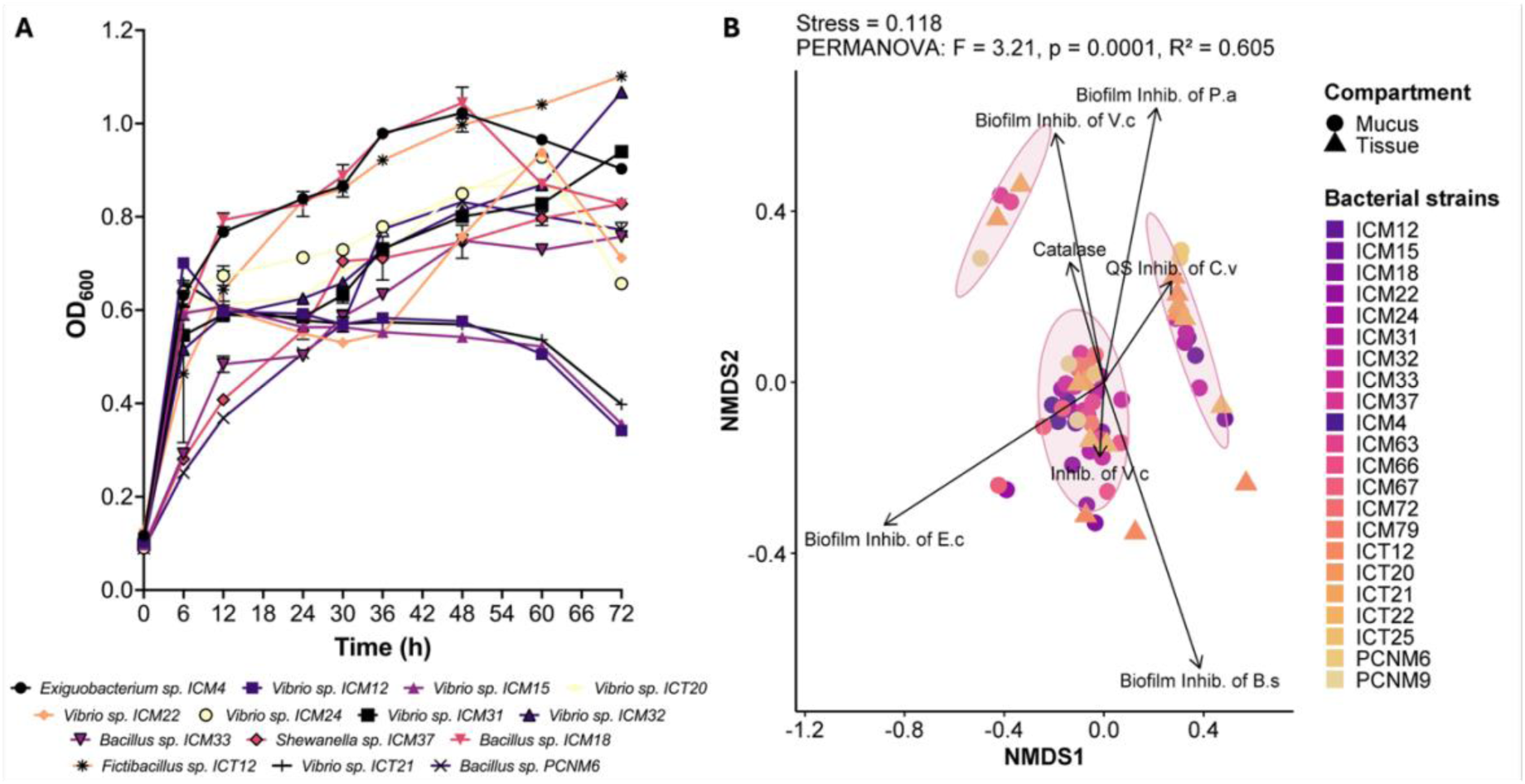
Growth performance and functional trait diversity among bacterial isolates from the urban coral *Madracis auretenra*. **(A)** Growth kinetics of 14 representative isolates evaluated over 72 h in liquid broth at 28°C. Optical density at 600 nm (OD_600_) was recorded at regular intervals. Most isolates reached stationary phase by 48–72 h, with *Exiguobacterium* sp. ICM4, *Bacillus* sp. ICM18, and *Fictibacillus* sp. ICT12 showing the highest OD_600_ values. Symbols represent mean values (n = 3) with error bars showing standard deviation. **(B)** Non-metric multidimensional scaling (NMDS) ordination plot based on Bray–Curtis similarity matrix of probiotic trait profiles (catalase activity, biofilm inhibition against four targets, quorum sensing inhibition, and pathogen antagonism). Stress = 0.118. Ellipses group isolates by coral compartment (mucus = circles; tissue = triangles). Vectors indicate trait contributions to observed clustering. PERMANOVA results (*F* = 3.21, p = 0.0001, R^2^ = 0.605) indicate significant differences among isolates based on functional profiles.

### Patterns in functional profiling of bacterial isolates

All isolates exhibited significantly different activity profiles when analyzed together, (PERMANOVA; *Pseudo-F* = 3.21; *p* = 0.0001; Table S6). The differences were graphed according to the post-hoc Monte Carlo analysis (Fig. S6), highlighting *Exiguobacterium* sp. ICM4, *Priestia* sp. ICM18, *Vibrio* sp. ICM67 and ICT20 as the bacteria that contributed the most to the differences found and indicating that some strains performed substantially better than others. SIMPER analysis (Table S7) revealed that *Exiguobacterium* sp. ICM4, *Priestia sp.* ICM18*, Bacillus* sp. ICM33, and *Vibrio* sp. ICM79 contributed disproportionately to the observed dissimilarity in biofilm inhibition activity, together accounting for over 90% of the variation compared to other isolates.

NMDS analysis revealed clear differentiation patterns in probiotic trait profiles among the bacterial isolates (stress = 0.118), with trait vectors explaining a substantial portion of the observed variation (R^2^ = 0.605). Strains were distributed along gradients of biofilm inhibition capacity, antagonistic activity, quorum sensing interference, and catalase production. Mucus-derived isolates formed a tighter cluster, while tissue-derived strains displayed greater functional divergence (Fig. 4B). Notably, *Fictibacillus* sp. ICT12, isolated from tissue, was clearly separated from the main cluster, reflecting a distinct functional profile. Both *Fictibacillus* sp. ICT12 and *Vibrio* sp. ICT21 were strongly associated with catalase activity, quorum sensing inhibition, and antagonism against *V. coralliilyticus*. In contrast, several mucus-derived strains—such as *Vibrio* sp. ICM12, *Bacillus* sp. ICM33, and PCNM6—occupied central positions in the NMDS plot, indicating functional redundancy across multiple activities. Collectively, these findings underscore the high antioxidant and pathogen-suppression potential of bacteria associated with the urban coral *Madracis auretenra* positioning them as promising candidates for inclusion in a coral probiotic consortium.

## Discussion

This study identified functionally diverse coral-associated bacteria from urban coral *Madracis auretenra* with traits supportive of probiotic potential, particularly for application in coral reef health management under stress conditions. Isolates affiliated with *Vibrio* were notable for consistently displaying multiple beneficial traits across all assays. While members of this genus are often excluded from probiotic consortia due to associations with coral disease (Rosado et al., 2019; Santoro et al., 2021), emerging evidence highlights their functional plasticity and ecological importance within the coral holobiont (Ma et al., 2018; Moynihan et al., 2022; Munn, 2015). Several *Vibrio* species contribute to coral fitness through roles such as nitrogen fixation in the mucus layer (Chimetto et al., 2008; Olson et al., 2009) and antimicrobial activity against pathogenic vibrios (Ritchie, 2006). In our previous work, we found *Vibrio* to be a dominant and stable component of the microbiome of healthy coral *M. auretenra*, with consistent populations across seasons, locations, and environmental stressors (Ruiz-Toquica et al., 2023; 2025). The identification of *Vibrio* strains in the present study with antioxidant activity, quorum sensing inhibition, and antibiofilm properties further supports a more nuanced understanding of their role within the coral microbiome and their potential inclusion in future probiotic consortia—an area that merits continued investigation.

Antioxidant production emerged as a widespread trait, with all isolates exhibiting catalase activity and several producing carotenoid-type pigments. Notably, *Vibrio* and *Shewanella* strains displayed catalase levels equal to or exceeding that of the positive control, suggesting a strong capacity to scavenge reactive oxygen species (ROS) (Dungan et al., 2021). These enzymatic defenses are central to coral resilience under thermal stress, mitigating oxidative damage that contributes to bleaching (Kusmita et al., 2023; Nielsen et al., 2018; Rosado et al., 2019). The detection of carotenoid-like compounds, based on spectrophotometric and chromatographic profiles, further supports the antioxidant potential of these bacteria (Vila et al., 2019). Although compound identity remains to be confirmed, UV-visible absorbance patterns indicated the presence of conjugated hydrophobic pigments such as zeaxanthins, β-carotenes, lycopenes, or astaxanthins, which are known to confer photoprotection and oxidative stress resistance (Reis-Mansur et al., 2019; Vila et al., 2019; Sajjad et al., 2020). Prior studies have shown that carotenoid accumulation within the coral holobiont provides a favorable biochemical environment to mitigate oxidative stress (Hillyer et al., 2017), reinforcing the relevance of these bacterial pigments and highlighting the need for further biochemical and functional characterization.

Pathogen control is another key probiotic function evaluated here. While direct antagonism against *Vibrio coralliilyticus* was relatively weak across isolates, biofilm inhibition and quorum-sensing (QS) disruption were widespread and strain-specific. Several isolates—particularly *Exiguobacterium* ICM4, *Bacillus* PCNM6, and *Vibrio* ICM67— exhibited robust inhibition of biofilms formed by both opportunistic and pathogenic bacteria. This suggests that suppression of pathogen colonization may occur through interference with communication systems rather than direct antibacterial activity (Gowrishankar et al., 2012; Ritchie, 2006). Supporting this, previous studies have documented the ability of marine bacteria, including *Vibrio* and *Bacillus*, to interfere with QS pathways and biofilm maturation (Irudayarajan et al., 2024; Rajasabapathy et al., 2020). In particular, the application of QS inhibitors has been shown to arrest the progression of coral disease by reducing the abundance of biofilm-forming opportunists (Certner & Vollmer, 2018), underscoring the potential of QS-disruptive and antibiofilm bacteria as biocontrol agents within coral holobionts. Several isolates inhibited violacein production in *Chromobacterium violaceum*, a model QS reporter. Although inhibition levels were moderate (< 40%) in liquid culture, differences based on the isolate’s origin (mucus vs. tissue) suggest microhabitat-specific functional profiles (Marchioro et al., 2020). Isolates such as *Fictibacillus* ICT12 and *Bacillus* PCNM6 were particularly effective in this regard, aligning with their broader antagonistic and antioxidant activity profiles.

The presence of untested genera—*Exiguobacterium*, *Fictibacillus*, and *Priestia*— within the high-performing isolates highlights the underexplored probiotic potential of bacteria beyond the frequently cited *Pseudoalteromonas* or *Cobetia* (Pitts et al., 2025; Rosado et al., 2019; Silva et al., 2021; Ushijima et al., 2023). These genera have demonstrated probiotic roles in other marine hosts, including oysters and shrimp (Gao et al., 2024; Kim et al., 2022; Zhang et al., 2024), and their inclusion here expands the phylogenetic and functional diversity of candidate coral probiotics. Notably, *Fictibacillus*, previously linked to protease production and nitrogen cycling in corals (Su et al., 2020), showed strong antioxidant and QS inhibition traits, supporting its potential role in holobiont homeostasis.

Beyond individual traits, the multidimensional functional profiles revealed clear differentiation among isolates with clusters corresponding to microhabitat origin, with mucus-derived strains showing greater functional overlap and tissue-derived isolates exhibiting more distinct profiles. *Fictibacillus* ICT12 stood out as an outlier, expressing multiple high-level traits and occupying a unique position in the multivariate trait space. These patterns suggest niche partitioning and specialization within the coral microbiome, likely shaped by local conditions and host interactions (Hernandez-Agreda et al., 2016; Ricci et al., 2019).

Importantly, several candidate strains exhibited functional redundancy, expressing multiple beneficial traits simultaneously. This is a desirable feature for probiotic consortia, as it enhances resilience and stability in the face of environmental variability or microbial turnover (Cárdenas et al., 2022; Peixoto et al., 2021). Unlike many coral probiotics assembled from single-trait strains (Rosado et al., 2019; Santoro et al., 2021), the isolates identified here offer overlapping antioxidant, QS-disruptive, and antibiofilm properties, supporting their integration into more robust consortia.

This work also evaluated practical considerations for probiotic selection, including stable growth performance and distinctive colony features (Peixoto et al., 2017; Villela et al., 2019). While some promising isolates were excluded due to inconsistent growth or close phylogenetic relationship with potentially coral pathogens or opportunists, the remaining candidates span five genera with complementary functions. This ecological and phenotypic diversity supports their future application in multi-strain formulations aimed at mitigating stress impacts on reef-building corals.

## Conclusions

The cultivable microbiota of the urban coral *Madracis auretenra* harbors functionally diverse bacteria exhibiting antioxidant and antibiofilm properties relevant to coral probiotics development. These findings underscore the ecological value of urban reef microbiomes as reservoirs of beneficial microbes and provide a foundation for developing next-generation probiotics for coral conservation. Future work should validate their effects *in vivo*, including testing in model systems and under stress conditions relevant to reef degradation.

## Supporting information

Supplementary Materia

## Acknowledgments

This study was supported by Internal Call #21 of 2021 from the Universidad de Bogotá Jorge Tadeo Lozano and by the “Alejandro Ángel Escobar” Foundation through the Colombia Biodiversa Fellowship (Call I-2023). We thank the Universidad Jorge Tadeo Lozano, Santa Marta campus, and the Water Quality Laboratory at the Universidad del Magdalena for providing laboratory facilities. We are also grateful to the “Ministerio de Ciencia y Tecnología” (Minciencias) for the Bicentennial Excellence Doctoral Fellowship, which supported part of this work. Special thanks to the research group “Estudio y Aprovechamiento de Productos Naturales Marinos y Frutas de Colombia” for their assistance with chromatographic analyses, and to Dr. Kim Ritchie for providing the *V. coralliilyticus* strain.

