## Supplementary Materia for "Functionally Diverse Coral-Associated Bacteria from an Urban Reef Exhibit Antioxidant and Antibiofilm Properties Relevant to Probiotic Development"

**Table S1.** General characteristics of bacterial isolates obtained from mucus and tissue of *Madracis auretenra* coral. Taken and modified from Ruiz-Toquica *et al.* (2023).

| Isolate ID | Source | Morphology | Gram | Taxon (genus) | Access code GenBank | Catalase | Pigments | Siderophores | Antagonism | AntiQS* | Quantitative characterization |
|---|---|---|---|---|---|---|---|---|---|---|---|
| **ICM 4** | Mucus | medium rod | +ve | *Exiguobacterium* | OL964632 | ++++ | Yes | No | No | Yes | Yes |
| **ICM12** | Mucus | curved rod | -ve | *Vibrio* | OL964633 | ++++ | Yes | Yes (Ca) | No | Yes | Yes |
| **ICM15** | Mucus | curved rod | -ve | *Vibrio* | OL964634 | ++++ | No | Yes (Ca) | No | Yes | Yes |
| **ICM18** | Mucus | long rod | +ve | *Priestia* | OP339931 | ++++ | Yes | Yes (Ca) | No | No | Yes |
| **ICM22** | Mucus | curved rod | -ve | *Vibrio* | OL964635 | ++++ | No | Yes (Ca) | No | Yes | Yes |
| **ICM24** | Mucus | curved rod | -ve | *Vibrio* | OL964636 | ++++ | No | Yes (Hy) | No | Yes | Yes |
| **ICM25** | Mucus | curved rod | -ve | *Vibrio* | OL964637 | ++++ | No | Yes (Hy) | No | Yes | Excluded |
| **ICM29** | Mucus | curved rod | -ve | *Vibrio* | OL964638 | ++++ | Yes | Yes (Ca) | No | No | Excluded |
| **ICM31** | Mucus | curved rod | -ve | *Vibrio* | OL964639 | ++++ | No | Yes (Hy) | No | Yes | Yes |
| **ICM32** | Mucus | curved rod | -ve | *Vibrio* | OL964640 | ++++ | No | Yes (Hy) | No | Yes | Yes |
| **ICM33** | Mucus | small rod | +ve | *Bacillus* | OL964641 | ++++ | Yes | Yes (Hy) | No | Yes | Yes |
| **ICM37** | Mucus | small rod | -ve | *Shewanella* | OL964642 | + | Yes | Yes (Hy) | No | Yes | Yes |
| **ICM39** | Mucus | curved rod long | -ve | *Vibrio* | OL964643 | +++ | No | Yes (Hy) | No | Yes | Excluded |
| **ICM63** | Mucus | small rod curved | -ve | *Vibrio* | OP339963 | ++++ | No | Yes | No | Yes | Yes |
| **ICM64** | Mucus | long rod | -ve | *Vibrio* | OP339964 | +++ | Yes | Yes | No | Yes | Excluded |
| **ICM66** | Mucus | small rod | -ve | *Vibrio* | OP339966 | ++++ | No | Yes | No | Yes | Yes |
| **ICM67** | Mucus | cocci-small rod | -ve | *Vibrio* | OP339967 | ++++ | No | Yes | No | Yes | Yes |
| **ICM72** | Mucus | small rod curved | -ve | *Vibrio* | OP339972 | +++ | No | Yes | No | Yes | Yes |
| **ICM79** | Mucus | small rod | -ve | *Vibrio* | OP339979 | ++++ | No | No | No | Yes | Yes |
| **ICT2** | Tissue | curved rod | -ve | *Vibrio* | OL964644 | ++++ | No | Yes (Hy) | No | Yes | Excluded |
| **ICT12** | Tissue | filamentous rod | +ve | *Fictibacillus* | OL964645 | ++++ | Yes | No | No | Yes | Yes |

| | | | | | | | | | | | |
|---|---|---|---|---|---|---|---|---|---|---|---|
| **ICT16** | Tissue | curved rod | -ve | *Vibrio* | OL964646 | ++++ | No | Yes (Hy) | No | Yes | Excluded |
| **ICT17** | Tissue | curved rod | -ve | *Vibrio* | OL964647 | No | No | Yes (Hy) | Yes* | Yes | Excluded |
| **ICT20** | Tissue | curved rod | -ve | *Vibrio* | OL964648 | ++++ | No | Yes (Hy) | No | Yes | Yes |
| **ICT21** | Tissue | curved rod | -ve | *Vibrio* | OL964649 | ++++ | No | Yes (Hy) | No | Yes | Yes |
| **ICT22** | Tissue | cocci-small rod | -ve | *Vibrio* | OP339997 | ++++ | No | Yes | No | Yes | Yes |
| **ICT24** | Tissue | cocci-small rod | -ve | *Vibrio* | OP339999 | ++++ | No | Yes | No | Yes | Excluded |
| **ICT25** | Tissue | small rod | -ve | *Vibrio* | OP340000 | ++++ | No | Yes | No | Yes | Yes |
| **ICT35** | Tissue | small rod | -ve | *Vibrio* | OP340010 | +++ | Yes | Yes | Yes | No | Excluded |
| **PCNM6** | Mucus | long sporulated rod (chain) | +ve | *Bacillus* | OL964650 | No | No | Yes (Ca) | Yes* | Yes | Yes |
| **PCNM9** | Mucus | medium rod | -ve | *Shewanella* | OL964651 | +++ | Yes | Yes (Hy) | No | Yes | Yes |
| **PCNM4** | Mucus | filamentous rod | +ve | *Nocardiopsis* | OL964652 | +++ | Yes | No | No | Yes | Excluded |

++++: very active; +++: active; *: against *Photobacterium* sp.; **Ca:** Carboxylates; **Hy:** Hydroxamates.

**Table S2.** Access codes of bacterial sequences of the genus *Vibrio*, isolated from healthy and diseased scleractinian corals, collected by Sweet *et al.* (2021) and from the present work (identified with the letters ICM/ICT).

| Access code GenBank | Taxon (genus) | Source | Condition |
|---|---|---|---|
| MK308630 | *(Vibrio parahemolyticus)* | Tissue | Healthy |
| MK308628 | *(Vibrio coralliilyticus)* | Tissue | Healthy |
| MK308627 | *(Vibrio alginolyticus)* | Tissue | Healthy |
| MK308612 | *(Vibrio owensii)* | Tissue | Healthy |
| MK308610 | *(Vibrio parahemolyticus)* | Tissue | Healthy |
| MK308601 | *(Vibrio coralliilyticus)* | Tissue | Healthy |
| MK308600 | *(Vibrio coralliilyticus)* | Tissue | Healthy |
| MK308595 | *(Vibrio coralliilyticus)* | Tissue | Healthy |
| MK308592 | *(Vibrio parahemolyticus)* | Tissue | Healthy |
| MK308588 | *(Vibrio diabolicus)* | Tissue | Healthy |
| MK308587 | *(Vibrio alginolyticus)* | Tissue | Healthy |
| MK308583 | *(Vibrio parahemolyticus)* | Tissue | Healthy |
| MK308581 | *(Vibrio alginolyticus)* | Tissue | Healthy |
| MK308579 | *(Vibrio parahemolyticus)* | Tissue | Healthy |
| JX221045 | (*Vibrio* sp.) | Tissue | Diseased |
| JX221044 | (*Vibrio* sp.) | Tissue | Diseased |
| JQ409387 | (*Vibrio* sp.) | Mucus | Healthy |
| JQ409386 | (*Vibrio* sp.) | Mucus | Healthy |
| JQ409385 | (*Vibrio* sp.) | Mucus | Healthy |
| JQ409384 | (*Vibrio* sp.) | Mucus | Healthy |
| JQ409383 | (*Vibrio* sp.) | Mucus | Healthy |
| JQ409380 | (*Vibrio* sp.) | Mucus | Healthy |
| JQ409374 | (*Vibrio* sp.) | Mucus | Healthy |
| JQ409373 | (*Vibrio* sp.) | Mucus | Healthy |
| JQ409372 | (*Vibrio* sp.) | Mucus | Healthy |
| JQ409371 | (*Vibrio* sp.) | Mucus | Healthy |
| JF411573 | (*Vibrio* sp.) | Tissue | Diseased |
| JF411571 | (*Vibrio* sp.) | Tissue | Diseased |
| JF411569 | (*Vibrio* sp.) | Tissue | Diseased |
| JF411562 | (*Vibrio* sp.) | Tissue | Diseased |
| JF411561 | (*Vibrio* sp.) | Tissue | Diseased |
| JF411560 | (*Vibrio* sp.) | Tissue | Diseased |
| JF411558 | (*Vibrio* sp.) | Tissue | Diseased |
| JF411557 | (*Vibrio* sp.) | Tissue | Diseased |
| JF411556 | (*Vibrio* sp.) | Tissue | Diseased |
| JF411555 | (*Vibrio* sp.) | Tissue | Diseased |
| JF411554 | (*Vibrio* sp.) | Tissue | Diseased |
| JF411553 | (*Vibrio* sp.) | Tissue | Diseased |

| | | | |
|---|---|---|---|
| JF411552 | (*Vibrio* sp.) | Tissue | Diseased |
| JF411551 | (*Vibrio* sp.) | Tissue | Diseased |
| JF411549 | (*Vibrio* sp.) | Tissue | Healthy |
| JF411543 | (*Vibrio* sp.) | Tissue | Healthy |
| JF411542 | (*Vibrio* sp.) | Tissue | Healthy |
| JF411541 | (*Vibrio* sp.) | Tissue | Healthy |
| JF411540 | (*Vibrio* sp.) | Tissue | Healthy |
| JF411539 | (*Vibrio* sp.) | Tissue | Healthy |
| JF411537 | (*Vibrio* sp.) | Tissue | Healthy |
| JF411536 | (*Vibrio* sp.) | Tissue | Healthy |
| JF411533 | (*Vibrio* sp.) | Tissue | Healthy |
| JF411532 | (*Vibrio* sp.) | Tissue | Healthy |
| JF411530 | (*Vibrio* sp.) | Tissue | Healthy |
| JF411529 | (*Vibrio* sp.) | Tissue | Healthy |
| JF411528 | (*Vibrio* sp.) | Tissue | Healthy |
| JF411526 | (*Vibrio* sp.) | Tissue | Diseased |
| JF411525 | (*Vibrio* sp.) | Tissue | Diseased |
| JF411522 | (*Vibrio* sp.) | Tissue | Diseased |
| JF411521 | (*Vibrio* sp.) | Tissue | Diseased |
| JF411518 | (*Vibrio* sp.) | Tissue | Diseased |
| JF411516 | (*Vibrio* sp.) | Tissue | Diseased |
| JF411509 | (*Vibrio* sp.) | Tissue | Diseased |
| JF411508 | (*Vibrio* sp.) | Tissue | Diseased |
| JF411482 | (*Vibrio* sp.) | Tissue | Healthy |
| JF411447 | (*Vibrio* sp.) | Tissue | Healthy |
| JF411443 | (*Vibrio* sp.) | Tissue | Healthy |
| JF411442 | (*Vibrio* sp.) | Tissue | Healthy |
| JF411438 | (*Vibrio* sp.) | Tissue | Diseased |
| JF411435 | (*Vibrio* sp.) | Tissue | Diseased |
| JF411433 | (*Vibrio* sp.) | Tissue | Diseased |
| JF411432 | (*Vibrio* sp.) | Tissue | Diseased |
| JF411425 | (*Vibrio* sp.) | Tissue | Diseased |
| JF411421 | (*Vibrio* sp.) | Tissue | Healthy |
| FJ952828 | (*Vibrio* sp.) | Tissue | Healthy |
| FJ952824 | (*Vibrio* sp.) | Tissue | Healthy |
| FJ952823 | (*Vibrio* sp.) | Tissue | Healthy |
| FJ952822 | (*Vibrio* sp.) | Tissue | Healthy |
| FJ952821 | (*Vibrio* sp.) | Tissue | Healthy |
| FJ952820 | (*Vibrio* sp.) | Tissue | Healthy |
| FJ952819 | (*Vibrio* sp.) | Tissue | Healthy |
| FJ952814 | (*Vibrio* sp.) | Tissue | Healthy |
| FJ952813 | (*Vibrio* sp.) | Tissue | Healthy |
| FJ952809 | (*Vibrio* sp.) | Tissue | Healthy |

| | | | |
|---|---|---|---|
| FJ952808 | (*Vibrio* sp.) | Tissue | Healthy |
| FJ952788 | (*Vibrio* sp.) | Tissue | Healthy |
| FJ952785 | (*Vibrio* sp.) | Tissue | Healthy |
| FJ952784 | (*Vibrio* sp.) | Tissue | Healthy |
| FJ952778 | (*Vibrio* sp.) | Tissue | Healthy |
| FJ952777 | (*Vibrio* sp.) | Tissue | Healthy |
| FJ952776 | (*Vibrio* sp.) | Tissue | Healthy |
| FJ952771 | (*Vibrio* sp.) | Tissue | Healthy |
| FJ952769 | (*Vibrio* sp.) | Tissue | Healthy |
| FJ952767 | (*Vibrio* sp.) | Tissue | Healthy |
| JF792070 | (*Vibrio* sp.) | Mucus | Healthy |
| JF792073 | (*Vibrio* sp.) | Mucus | Healthy |
| OL964633 | (ICM12) | Mucus | Healthy |
| OL964634 | (ICM15) | Mucus | Healthy |
| OL964635 | (ICM22) | Mucus | Healthy |
| OL964636 | (ICM24) | Mucus | Healthy |
| OL964637 | (ICM25) | Mucus | Healthy |
| OL964638 | (ICM29) | Mucus | Healthy |
| OL964639 | (ICM31) | Mucus | Healthy |
| OL964640 | (ICM32) | Mucus | Healthy |
| OL964643 | (ICM39) | Mucus | Healthy |
| OP339963 | (ICM63) | Mucus | Healthy |
| OP339964 | (ICM64) | Mucus | Healthy |
| OP339966 | (ICM66) | Mucus | Healthy |
| OP339967 | (ICM67) | Mucus | Healthy |
| OP339972 | (ICM72) | Mucus | Healthy |
| OP339979 | (ICM79) | Mucus | Healthy |
| OL964644 | (ICT2) | Tissue | Healthy |
| OL964646 | (ICT16) | Tissue | Healthy |
| OL964647 | (ICT17) | Tissue | Healthy |
| OL964648 | (ICT20) | Tissue | Healthy |
| OL964649 | (ICT21) | Tissue | Healthy |
| OP339997 | (ICT22) | Tissue | Healthy |
| OP339999 | (ICT24) | Tissue | Healthy |
| OP340000 | (ICT25) | Tissue | Healthy |
| OP340010 | (ICT35) | Tissue | Healthy |
| NR_102775.2 | *(Thermotoga maritima)* | | |

**Table S3.** Probiotic characteristics of bacterial isolates from *M. auretenra* coral. According to spectrophotometric results.

| Isolate ID | Taxon (genus) | Access code | Relationship with pathogens | Catalase | Anti-QS (Biofilms) | Anti-QS (*C. violaceum*) | Antagonism (*V. coralliilyticus)* |
|---|---|---|---|---|---|---|---|
| ICM4 | *Exiguobacterium* | OL964632 | N.A. | Regular | High | Low | Null |
| ICM12 | *Vibrio* | OL964633 | No | High | High | Low | Null |
| ICM15 | *Vibrio* | OL964634 | No | Low | High | Low | Low |
| ICM18 | *Priestia* | OP339931 | N.A. | Regular | High | Low | Low |
| ICM22 | *Vibrio* | OL964635 | No | High | High | Low | Null |
| ICM24 | *Vibrio* | OL964636 | No | Regular | High | Null | Low |
| ICM31 | *Vibrio* | OL964639 | No | High | High | Low | Low |
| ICM32 | *Vibrio* | OL964640 | No | High | High | Null | Low |
| ICM33 | *Bacillus* | OL964641 | N.A. | High | High | Null | Low |
| ICM37 | *Shewanella* | OL964642 | N.A. | High | High | Low | Low |
| ICM63 | *Vibrio* | OP339963 | No | Low | High | Low | Low |
| ICM66 | *Vibrio* | OP339966 | No | Regular | High | Low | Low |
| ICM67 | *Vibrio* | OP339967 | No | Regular | High | Low | Low |
| ICM72 | *Vibrio* | OP339972 | No | High | High | Low | Low |
| ICM79 | *Vibrio* | OP339979 | No | High | High | Low | Low |
| ICT12 | *Fictibacillus* | OL964645 | N.A. | Regular | Null (*VC*) | Regular | Low |
| ICT20 | *Vibrio* | OL964648 | No | High | Null (*EC*) | Low | Low |
| ICT21 | *Vibrio* | OL964649 | No | High | High | Low | Low |
| ICT22 | *Vibrio* | OP339997 | No | High | High | Low | Low |
| ICT25 | *Vibrio* | OP340000 | No | High | High | Low | Low |
| PCNM6 | *Bacillus* | OL964650 | N.A. | Regular | High | Regular | Low |
| PCNM9 | *Shewanella* | OL964651 | N.A. | Regular | High | Low | Low |

Isolates were named by collection site: Inca-Inca (**IC**) and Punta Cabeza de Negro (**PCN**); as well as source of extraction: mucus (**M**) and tissue (**T**); ***VC***: *V. coralliilyticus;* ***EC***: *E. coli.;* **N. A.:** Not applicable.

**Table S4.** Absorption peaks of pigments detected from UHPLC analysis.

| Isolate | Pigment color | Absorption peaks (nm) | Potential pigment type |
|---|---|---|---|
| *Exiguobacterium* sp. ICM4 | Orange | 465, 464, 445, 473, 465, 464 | β-Carotene, Zeaxanthin, β-Zeacarotene |
| *Vibrio* sp. ICM12 | Yellow, brown | 463, 463, 444, 466, 450, 463, 465 | β-Zeacarotene, Lycopene, Zeaxanthin |
| *Bacillus* sp. ICM33 | Pink, red | 482, 491, 493, 465, 455, 463 | β-Carotene, Canthaxanthin, Astaxanthin |

**Table S5.** Results of the statistical analysis applied to each of the probiotic traits evaluated.

| | | | | | |
|---|---|---|---|---|---|
| **one-way ANOVA** | | | | | |
| | **df** | **Sum Sq** | **Mean Sq** | **F value** | **Pr (>F)** |
| **Source** | 3 | 15,92 | 5,308 | 1,846 | **0,147** |
| **Residuals** | 68 | 195,54 | 2,876 | | |
| **Kruskal-wallis** | | | | | |
| **chi-squared** | | | | 185,33 | |
| **df** | | | | 3 | |
| **p-value** | | | | **<2,2E-16** | |
| **Mann-Whitney U** | | | | | |
| **W** | | | | 96 | |
| **p-value** | | | | **0,03954** | |
| **Welch two sample t-test** | | | | | |
| **t** | | | | -1,1718 | |
| **df** | | | | 15,725 | |
| **p-value** | | | | **0,2587** | |
| **95 % confidence interval** | | | | -5,238514 | |
| | | | | 1,512278 | |

**df:** degrees of freedom

**Table S6.** Results of PERMANOVA analysis of *M. auretenra* bacterial isolates based on tested probiotic activities.

| | df | SS | MS | Pseudo-F | P(perm) |
|---|---|---|---|---|---|
| Bacterial isolates | 21 | 11717 | 557,95 | 3,2077 | **0,0001** |
| Residual | 44 | 7653,4 | 173,94 | | |
| Total | 65 | 19370 | | | |

**df:** degrees of freedom; **SS:** sum of squares; **MS:** mean of squares; **Pseudo-F:** F statistic; **P(perm):** significance level.

**Table S7.** Similarity Percentage Analysis (SIMPER). Summary of the percentage contribution of biofilm inhibition to dissimilarity between isolates.

| Isolate ID | Avarage similarity | Contrib% biofilms |
|---|---|---|
| ICM4 | 94.88 | 83.33 |
| ICM12 | 81.68 | 72.83 |
| ICM15 | 80.74 | 96.68 |
| ICM18 | 92.87 | 72.81 |
| ICM22 | 86.75 | 74.93 |
| ICM24 | 80.99 | 82.16 |
| ICM31 | 85.38 | 79.39 |
| ICM32 | 85.51 | 76.63 |
| ICM33 | 93.71 | 86.09 |
| ICM37 | 82.60 | 82.23 |
| ICM63 | 78.49 | 72.37 |
| ICM66 | 88.53 | 92.78 |
| ICM67 | 88.86 | 75.06 |
| ICM72 | 85.34 | 83.72 |
| ICM79 | 91.19 | 87.29 |
| ICT12 | 74.04 | 80.33 |
| ICT20 | 89.91 | 73.08 |
| ICT21 | 71.69 | 83.53 |
| ICT22 | 69.52 | 81.56 |
| ICT25 | 78.81 | 81.73 |
| PCNM6 | 83.26 | 72.53 |
| PCNM9 | 79.81 | 84.69 |

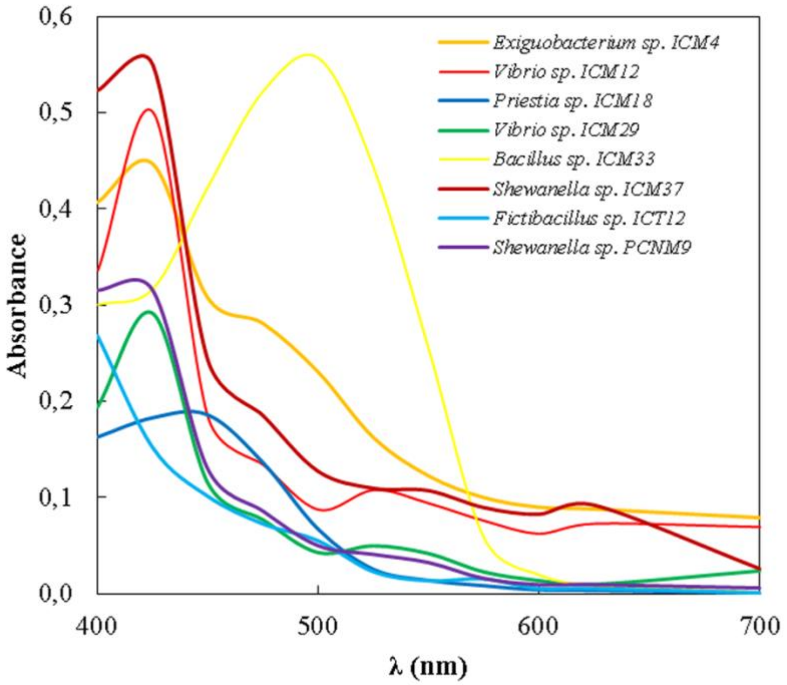


**Figure S1.** Absorption curves of pigments produced by isolates with initial probiotic potential.

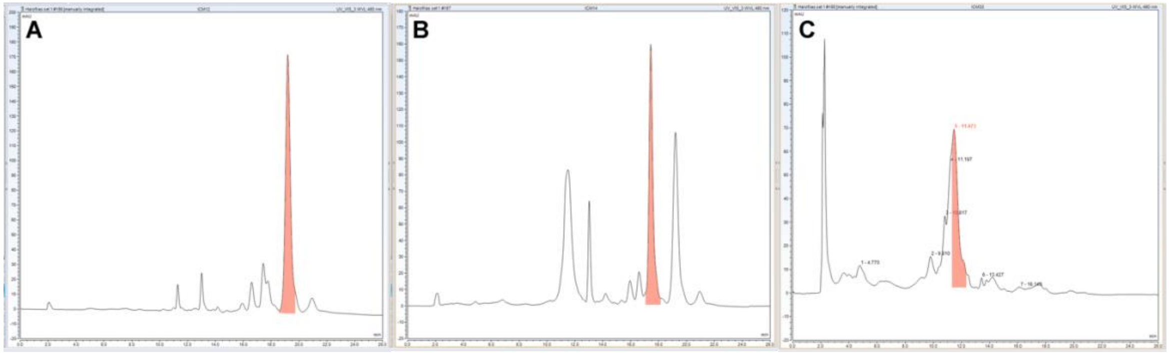


**Figure S2.** Chromatographic analysis of extracts from three bacterial isolates. **A)** ICM12; **B)** ICM4; **C)** ICM33.

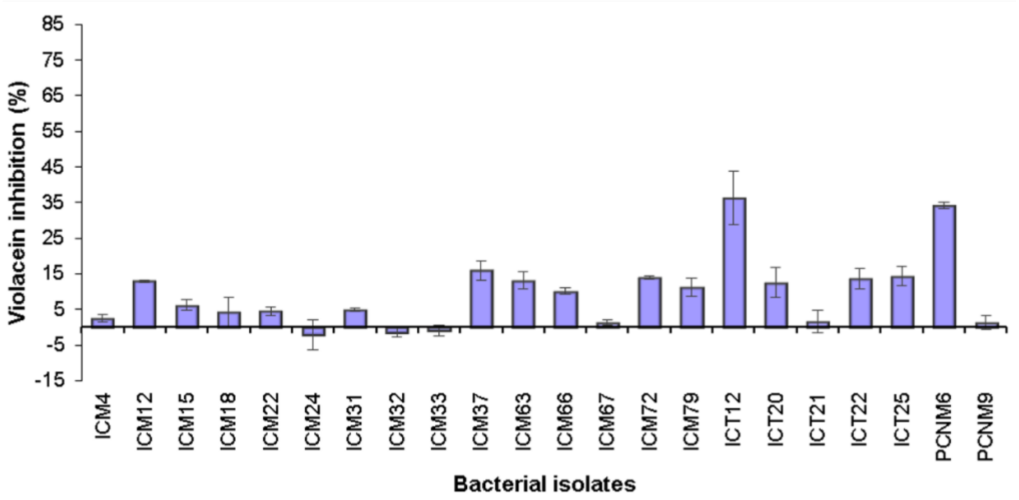


**Figure S3.** Inhibition of violacein pigment by bacterial isolates with probiotic potential. Data are presented as the mean ± SE (n = 3).

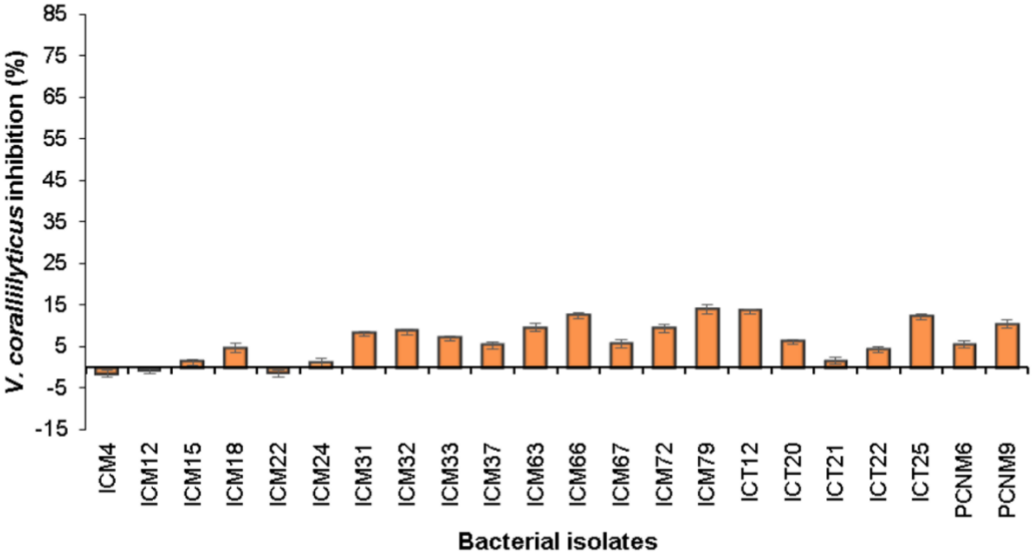


**Figure S4.** Antagonistic activity. Percentage inhibition of the coral pathogen *Vibrio coralliilyticus* by bacterial isolates with probiotic potential. Data are presented as the mean ± SE (n = 3).

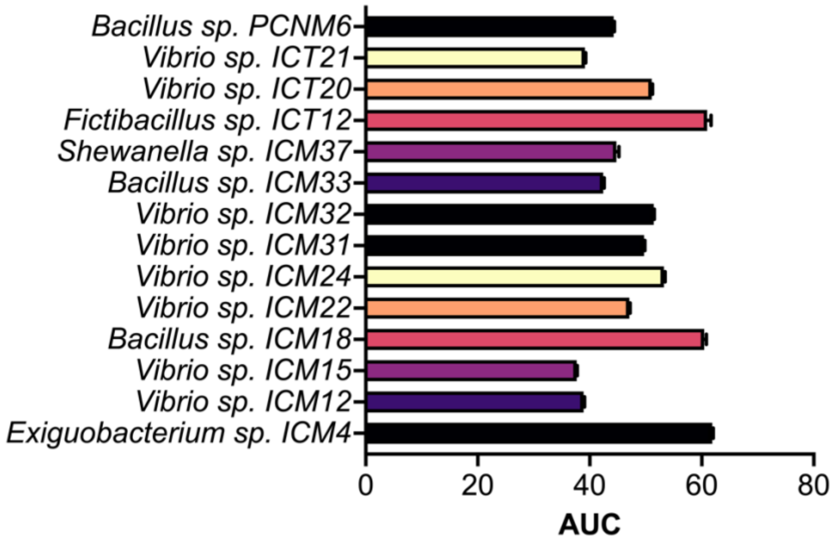


**Figure S5.** Area under the curve of bacterial isolates with probiotic potential.

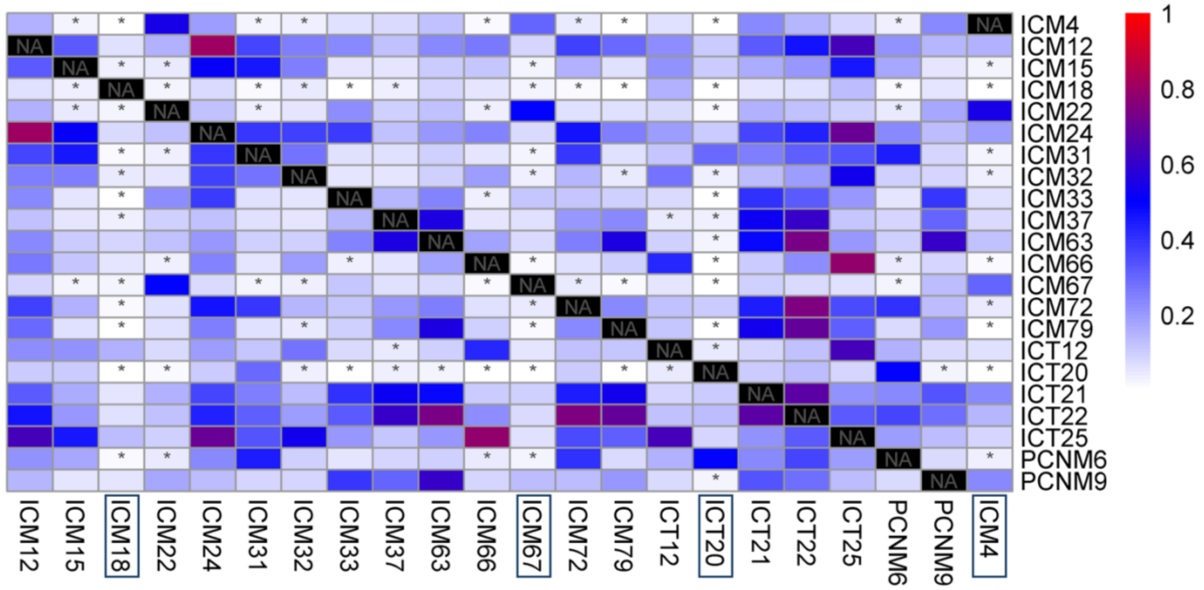


**Figure S6.** Monte Carlo pairwise test p-value heat map. Asterisks indicate significance among bacteria (* = $p < 0.05$).

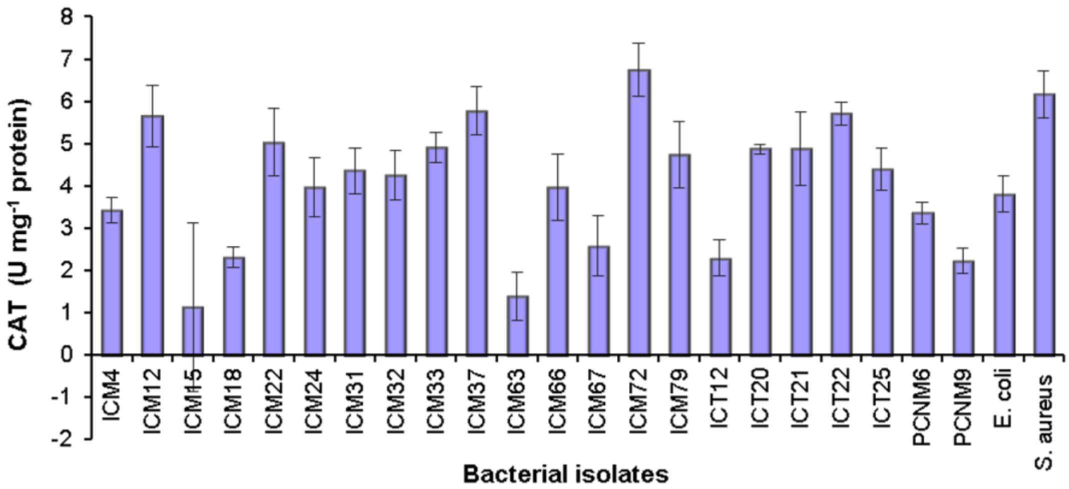


**Figure S7.** Catalase activity of bacterial isolates with probiotic potential. Data are presented as the mean ± SE (n = 3).

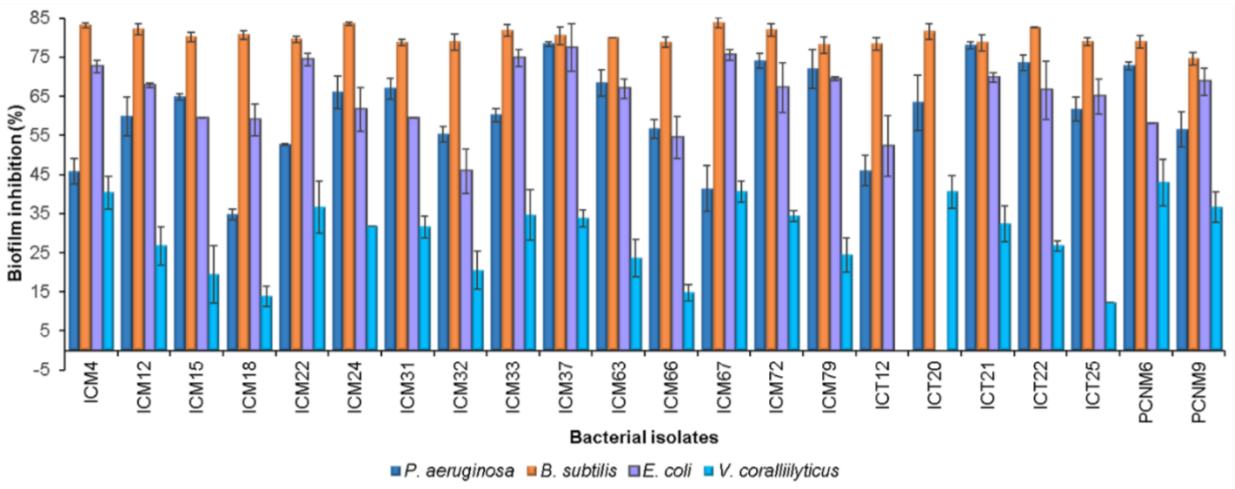


**Figure S8.** Inhibition of biofilm formation of four reporter strains (*Pseudomonas aeruginosa, Bacillus subtilis, Escherichia coli, Vibrio coralliilyticus*) by bacterial isolates with probiotic potential. Data are presented as the mean ± SE (n = 3).
